# Exo-LAMP: A Rapid Extraction-Free Isothermal Assay for Exosomal mRNA Detection

**DOI:** 10.64898/2026.09.18.752807

**Authors:** Md Akeruzzaman Shaon, Farjana Haque, Milkiyas Toru Tantu, Omar Hamza Bin Manjur, Doaa M. Hanafy, Sonia Tamanna, Kevin M. Koo, Allen G. Ross, Kiran Shrestha, Fatema Zerin Farhana, Muhammad J. A. Shiddiky

## Abstract

Preeclampsia arises from early placental dysfunction that precedes clinical presentation, creating a need for molecular readouts that enable timely risk assessment. Placenta derived exosomes carry nucleic acid biomarkers with diagnostic potential, yet current analytical workflows rely on multistep isolation and extraction procedures that are slow and resource intensive. Here, we report Exo-LAMP, an extraction-free isothermal assay for detecting placental exosomal *KISS1* mRNA, representing the first isothermal approach for exosomal mRNA detection. Placental alkaline phosphatase (PLAP) positive syncytiotrophoblast derived exosomes are selectively captured from biofluids using PLAP antibody conjugated magnetic beads within 20 minutes. Captured exosomes are treated with RNase inhibitor and lysed at 90 °C for 5 minutes, enabling direct RNA release while preserving transcript integrity and eliminating conventional extraction steps. The released RNA is analyzed in the same vessel by colorimetric reverse transcription loop-mediated isothermal amplification (RT-LAMP), providing semi-quantitative readout within 25 to 30 minutes. The assay quantifies *KISS1* mRNA across 10^3^ to 10^8^ exosomes per reaction, with an estimated limit of detection of about 2 exosomes and a limit of quantification of 10^3^ exosomes. Stoichiometric analysis indicates approximately one *KISS1* transcript per 27 exosomes, compared with one per 38 exosomes measured by reverse transcription quantitative polymerase chain reaction (RT-qPCR). In both the exosome-depleted serum spike-in and the plasma spike-in samples containing mixed vesicle populations, the assay detects target derived transcripts at 1% abundance and retains measurable signal at the quantification limit in the presence of up to 10^8^ non-target exosomes. These results establish Exo-LAMP as a rapid and simplified platform for exosomal mRNA analysis that integrates selective capture, lysis, and amplification in a single workflow completed in less than 1 hour. The method provides a general framework for semi-quantitative detection of exosomal transcripts and can be adapted to other coding and non-coding RNA targets for liquid biopsy applications.

**TABLE OF CONTENT:** TOC Figure

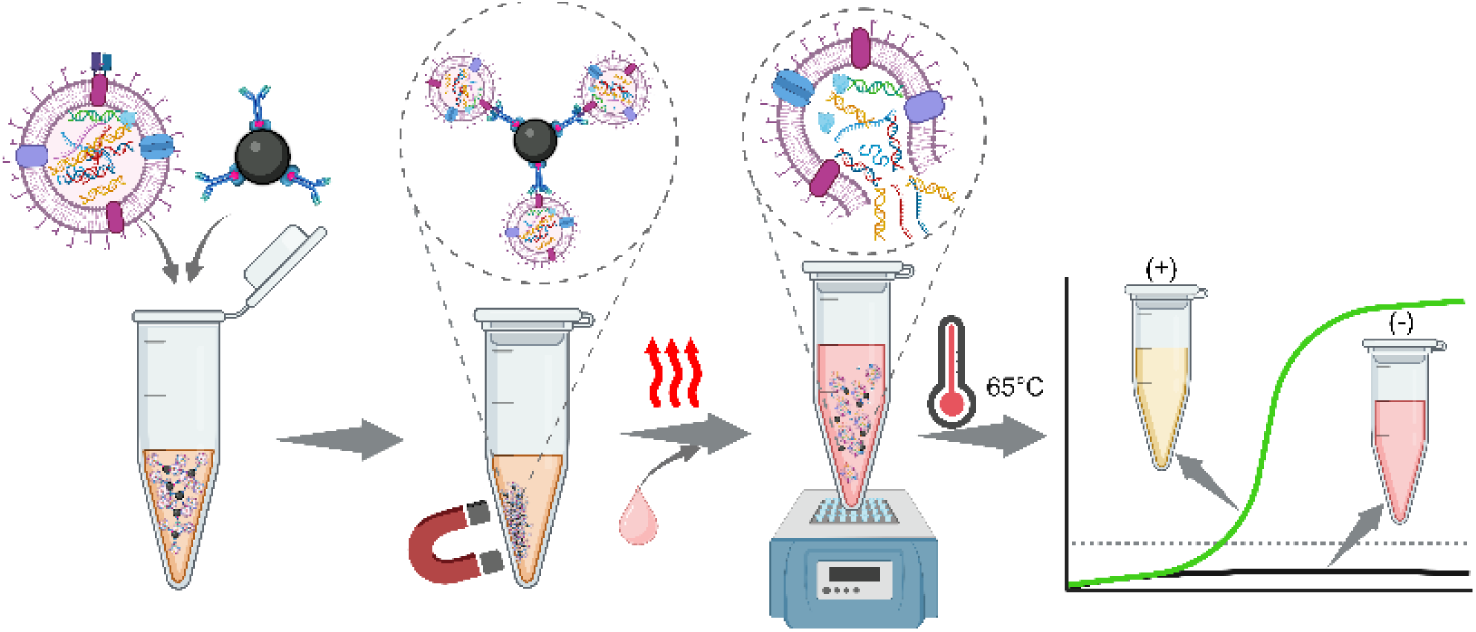

## 1. INTRODUCTION

Preeclampsia (PE) is a multisystem hypertensive disorder of pregnancy associated disorder defined by new onset hypertension (≥140/90 mmHg) with proteinuria (≥0.3 g/24 h), or, in the absence of proteinuria after 20 weeks of gestation [1, 2]. It affects 3 to 8% of pregnancies worldwide [3] and contributes to approximately 46,000 maternal and 500,000 fetal and neonatal deaths annually [4], with disproportionate impact in low-resource settings where access to early diagnostic testing and timely intervention is limited. The condition originates from abnormal placentation, where defective trophoblast invasion and incomplete spiral artery remodeling lead to placental hypoxia, oxidative stress, and release of placental factors into the maternal circulation [5]. These molecular changes arise before maternal symptoms become evident, which limits early diagnosis and timely intervention.

In this context, placenta derived extracellular vesicles (EVs), particularly exosomes, provide a direct molecular representation of placental function and can be detected in maternal blood from early gestation, offering access to disease related signals prior to clinical presentation [6]. Among their molecular cargo, KISS1 mRNA encodes kisspeptin peptides, a regulator of trophoblast invasion and placental development. Altered KISS1 expression has been linked to impaired trophoblast migration and shallow placentation in PE [7]. The lipid bilayer of exosomes protects RNA from RNase degradation [8], supporting their use as stable and minimally invasive biomarkers for monitoring placental health.

Despite its diagnostic relevance, exosomal mRNA analysis remains limited by complex multistep workflows that constrain analytical performance and clinical translation. Conventional strategies typically require sequential exosome isolation, RNA extraction, and downstream molecular detection [9]. Common isolation approaches, including ultracentrifugation, differential centrifugation, and commercial precipitation-based kits, are time intensive, labor demanding, and dependent on specialized instrumentation [9–11]. In addition, these methods generally enrich bulk extracellular vesicle populations rather than placental specific subtypes, which can dilute disease relevant signals in maternal circulation and reduce analytical specificity for early placental dysfunction.

Although electrochemical and microfluidic platforms have improved selectivity and sensitivity, many still depend on complex surface modification, enzymatic labeling, or sequential enrichment steps that increase procedural complexity and operational cost while limiting throughput [12–17]. To address these constraints, direct immunoaffinity capture based on a single placental surface marker offers a more streamlined analytical strategy. EVs retain membrane proteins reflective of their cellular origin [18], enabling targeted recognition of syncytiotrophoblast derived vesicles through placental alkaline phosphatase (PLAP) [19]. Anti-PLAP functionalized magnetic particles enable selective enrichment of PLAP positive (PLAP^P^) exosomes while minimizing interference from non-placental vesicle populations and removing the need for prior bulk isolation, thereby simplifying sample preparation and improving the specificity of downstream molecular analysis [20, 21].

Following selective capture, access to exosomal RNA remains a critical step that determines analytical performance. Conventional extraction strategies based on phenol chloroform partitioning or silica membrane purification introduce multiple handling steps, extend assay time, and increase the risk of sample loss, particularly when transcript abundance is low [22]. Heat-mediated lysis at 90 - 95 °C offers a simplified approach for releasing nucleic acids directly from vesicles without conventional extraction kit-based procedures [23, 24]. In addition, high-temperature treatment can substantially reduce nuclease activity, thereby helping to limit enzymatic degradation of released RNA [25]. However, RNA released during the heating transition may remain susceptible to RNase-mediated degradation before nuclease activity is sufficiently reduced, particularly in complex biological matrices [8]. Therefore, the inclusion of RNase inhibitors can provide additional protection for released RNA, helping to preserve RNA integrity and improve the reliability of downstream detection [26].

In contrast to exosomal protein and microRNA analysis, targeted detection of exosomal mRNA remains comparatively underdeveloped despite its relevance for functional readouts of gene expression. Current analytical strategies rely primarily on reverse transcription quantitative polymerase chain reaction (RT-qPCR), droplet digital PCR, or sequencing based workflows [1, 10, 26, 27]. While these methods provide high analytical sensitivity, they depend on purified RNA inputs, thermocycling equipment, and multistep processing that restricts adaptability for streamlined workflows. Isothermal amplification offers an alternative route for nucleic acid detection by eliminating thermal cycling requirements [28, 29]. In particular, reverse transcription loop-mediated isothermal amplification (RT-LAMP) enables rapid and specific target amplification at a constant temperature through the coordinated activity of multiple primers recognizing distinct regions of the sequence. This approach is compatible with crude lysates and supports direct signal generation without reliance on complex instrumentation, thereby reducing procedural complexity while maintaining analytical performance [28, 30].

Here, we report Exo-LAMP, an integrated workflow for semi-quantitative analysis of exosomal KISS1 mRNA that combines selective vesicle capture, RNA release, and nucleic acid detection within a single streamlined process; to our knowledge, Exo-LAMP represents the first isothermal method developed for exosomal mRNA detection. PLAP^P^ exosomes are directly isolated using PLAP antibody conjugated magnetic beads, followed by heat mediated lysis in the presence of an RNase inhibitor to enable extraction free release of RNA, and subsequent colorimetric RT-LAMP performed in the same reaction vessel. Using PLAP^P^ BeWo cells as a placental model and PLAP negative (PLAP^N^) HTR-8 cells as a negative control, the system demonstrates selective enrichment of placental vesicles while avoiding nonspecific bulk extracellular vesicle isolation. This design eliminates conventional RNA extraction and thermocycling requirements, thereby reducing procedural complexity and handling induced variability. The integrated workflow provides controlled access to exosomal RNA and enables direct isothermal detection under uniform reaction conditions, establishing a simplified analytical framework for exosomal mRNA measurement. This approach supports practical assessment of circulating placental signals and provides a basis for streamlined molecular evaluation of placental status for early risk stratification of PE.

## 2. MATERIALS AND METHODS

### 2.1 Cell culture and genomic DNA extraction

Placental alkaline phosphatase-positive (PLAP^P^) BeWo (CCL-98) human choriocarcinoma trophoblast cells and placental alkaline phosphatase–negative (PLAP^N^) HTR-8/SVneo (CRL-3271) human trophoblast cells were obtained from the American Type Culture Collection (ATCC, USA). Both cell lines were maintained in T-25 culture flasks (Life Technologies, Australia) at 37 °C in a humidified incubator containing 5% CO_2_. BeWo cells were cultured in Ham’s F-12K (Kaighn’s) medium, while HTR-8/SVneo cells were cultured in RPMI-1640 medium, each supplemented with 10% exosome-depleted fetal bovine serum and 1% penicillin-streptomycin (Life Technologies, Australia). After 60 h of incubation, the conditioned cell culture medium (CCM) containing secreted exosomes was collected for downstream analysis. Genomic DNA was extracted from the cell lines using the AllPrep DNA/RNA/miRNA Universal Kit (Qiagen GmbH, Germany) according to the manufacturer’s protocol. The concentration and purity of the extracted DNA was evaluated using a NanoDrop One microvolume UV-Vis spectrophotometer (Thermo Fisher Scientific).

### 2.2 Commercial kit-based exosome isolation and characterization

Exosomes were isolated from CCM supernatants using a centrifugation-based approach. First, the collected supernatant was centrifuged at 2,000 x g for 30 minutes to remove cells and large debris. This pre-cleared supernatant was then passed through a 0.22 µm syringe filter to eliminate remaining particles. Exosomes were subsequently concentrated using a Total Exosome Isolation Kit (Thermo Fisher Scientific, USA) according to the manufacturer’s instructions. Finally, the resulting exosome pellet was resuspended in 100 µL of nuclease-free phosphate buffered saline (PBS).

Nanoparticle tracking analysis (NTA) was performed using a NanoSight NS300 system (Malvern, PA, USA) to characterize the isolated EVs. Prior to analysis, the samples were diluted 100-fold with PBS to obtain an optimal particle concentration. The diluted samples were then introduced into the instrument, and measurements were conducted using the automated acquisition sequence: PUMPLOAD, REPEATSTART, PRIME, DELAY 10, CAPTURE 30, and REPEAT 5. Videos were recorded using a camera level of 13, shutter speed of 20 ms, and camera gain of 1500, and these parameters were kept constant for all samples to ensure consistency. The recorded videos were subsequently analyzed using the NanoSight software to calculate the particle concentration as well as the mean size, mode size, and overall size distribution of EVs.

### 2.3 Western blot analysis of exosomal PLAP

Western blotting was performed to confirm the presence of PLAP in exosomes. Briefly, isolated exosomes from PLAP^P^ BeWo and PLAP^N^ HTR-8 cells were mixed with sample buffer, heated at 95 °C for 5 min, separated by SDS-PAGE, and transferred onto a polyvinylidene difluoride (PVDF) membrane using the Trans-Blot Turbo transfer system (Bio-Rad). The membrane was then blocked and incubated with a primary anti-PLAP antibody, followed by a horseradish peroxidase (HRP)-conjugated secondary antibody. Protein bands were visualized using chemiluminescent substrate. All experiments were performed in duplicate.

### 2.4 Immunomagnetic isolation of exosomes and evaluation of capture efficiency

Anti-PLAP antibodies (ab218848, Abcam) were biotinylated using a Type B biotin conjugation kit (ab201796, Abcam) according to the manufacturer’s instructions. Briefly, 1 µL of modifier reagent was mixed with 10 µL of antibody solution (1 mg/mL). The mixture was then added to the lyophilized biotin reagent and gently resuspended by pipetting. The reaction was incubated in the dark at room temperature for 20 min to allow antibody–biotin conjugation. After incubation, 1 µL of quencher reagent was added, mixed gently, and the biotinylated antibodies were stored at 4 °C until further use. For antibody immobilization, 20 µL of Dynabeads Streptavidin MyOne magnetic beads (MB) (10 mg/mL) were washed twice with PBS and resuspended in 50 µL of biotinylated PLAP antibody solution (160 ng/mL). The mixture was incubated for 30 min at room temperature with gentle rocking to allow antibody binding to the beads. The antibody-functionalized beads (MB@Ab) were then washed three times with PBS to remove unbound antibodies and finally resuspended in 100 µL PBS and stored at 4 °C. For exosome capture, pre-cleared CCM was incubated with MB@Ab at a 20:1 volume ratio (100 µL CCM : 5 µL MB@Ab) for 30 min at room temperature with gentle rocking. This ratio was maintained to enable efficient capture of exosomes from larger CCM volumes. After incubation, the exosome-bound beads were washed three times with PBS to remove unbound components and finally resuspended in 20 µL PBS or exosome-depleted serum for further analysis.

To evaluate the efficiency of immunomagnetic separation, a known number of exosomes was spiked into fresh cell culture medium and exosome depleted serum samples. The samples were then incubated with the MB@Ab complex for 20 min at room temperature with gentle rocking, maintaining the optimized bead-to-sample ratio. After incubation, the mixture was centrifuged at 10,000 × g for 5 min, and the supernatant containing unbound EVs was collected. The number of remaining EVs in the supernatant was measured using NTA. Capture efficiency was calculated using the following equation:

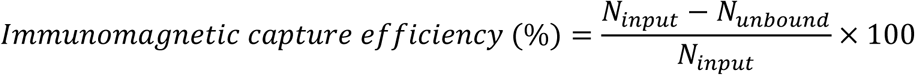

Where, *N_input_* = total number of exosomes initially added; *N_unbound_* = number of exosomes remaining in the supernatant after capture.

### 2.5 Zeta potential analysis

Zeta potential measurements were performed using a NANOTRAC WAVE II instrument (Microtrac MRB, USA). Ultrapure water was used as the dispersant with a refractive index of 1.333 and viscosity of 0.8872 mPa·s at 25 °C. The refractive index of extracellular vesicles was set to 1.37, while the refractive index of the magnetic beads was set to 1.59 based on typical values reported for lipid vesicles and polymer microspheres. For antibody-coated bead-EV complexes, an effective refractive index of approximately 1.45-1.50 was used to account for the composite structure of the particle surface.

### 2.6 Exosomal RNA extraction using a commercial kit and direct heat-mediated lysis

Exosomal total RNA was extracted from isolated exosomes using the Total Exosome RNA and Protein Isolation Kit (Thermo Fisher Scientific). Briefly, the aqueous phase obtained after acid-phenol:chloroform separation was mixed with 1.25 volumes of 100% ethanol and transferred to a silica-based filter cartridge to allow RNA binding. The bound RNA was sequentially washed with miRNA Wash Solution 1, followed by two washes with Wash Solution 2/3 to remove impurities. After an additional centrifugation step to remove residual wash buffer, RNA was eluted from the membrane using preheated elution buffer or nuclease-free water in two sequential 50 µL elution steps. The purified exosomal RNA was immediately placed on ice and stored at −20 °C until further analysis.

In parallel, an extraction-free direct heat-lysis approach was used to release exosomal nucleic acids. Briefly, immunomagnetically captured exosomes were subjected to thermal lysis at 90 °C and 95 °C, and for each temperature both 5 min and 10 min incubation times were tested, to determine the optimal temperature and time for efficient vesicle disruption. This heat treatment disrupts the exosomal lipid membrane and releases the total nucleic acid into solution. The resulting lysate was directly used as the template for RT-LAMP amplification without further RNA purification.

### 2.7 *KISS1* mRNA synthesis from gBlocks Gene Fragments

mRNA standards were synthesized from gBlocks Gene Fragments (Integrated DNA Technologies, IDT) using the HiScribe™ T7 Quick High Yield RNA Synthesis Kit (E2050S; New England Biolabs, NEB) according to the manufacturer’s instructions. For each transcription reaction (20 µL total volume), 10 µL of gBlocks template DNA (10 ng/µL) was combined with 8 µL of NTP mix and 2 µL of T7 RNA polymerase mix. The reaction mixture was incubated at 37 °C for 2 h to allow in vitro transcription of mRNA. Following transcription, the reaction was diluted with 30 µL of nuclease-free water. To remove residual template DNA, 2 µL of DNase I (2 U/µL) was added, and the mixture was incubated at 37 °C for 15 min. The synthesized mRNA was then purified using a lithium chloride (LiCl) precipitation method. Briefly, 25 µL of 7.5 M LiCl was added to each reaction, followed by incubation at −20 °C for 30 min to precipitate the RNA. The RNA pellet was collected by centrifugation at 17,000 ×g for 15 min at 4 °C. The pellet was washed with 500 µL of ice-cold 70% ethanol and then resuspended in 50 µL of nuclease-free water. The purified mRNA was serially diluted to generate standard solutions and stored at −20 °C until use. RNA concentration was determined using the NanoDrop One microvolume UV-Vis spectrophotometer.

### 2.8 Primer designing

A set of six loop-mediated isothermal amplification (LAMP) primers (FIP, BIP, F3, B3, LF, and LB) targeting KISS1 mRNA (NCBI accession ID: NM_002256.4) was designed using PrimerExplorer v5 (https://primerexplorer.eiken.co.jp/lampv5/index.html). In addition, two primers for reverse transcription quantitative polymerase chain reaction (RT-qPCR) (FP and RP) were designed using NCBI Primer-BLAST (https://www.ncbi.nlm.nih.gov/tools/primer-blast). All primers were evaluated for potential secondary structures, including hairpin formation, self-dimerization, and cross-dimerization, using the IDT OligoAnalyzer tool. To ensure selective amplification of KISS1 mRNA and to avoid amplification from genomic KISS1 DNA, the primer design targeted exon–exon junction regions of the transcript. This strategy preferentially amplifies spliced mRNA transcripts rather than genomic sequences containing introns. The specificity of each primer was further verified using BLASTn analysis against the NCBI human genome database to confirm the absence of significant off-target complementarity. The sequences of all primers used in this study are provided in Table S1.

**Table S1.**
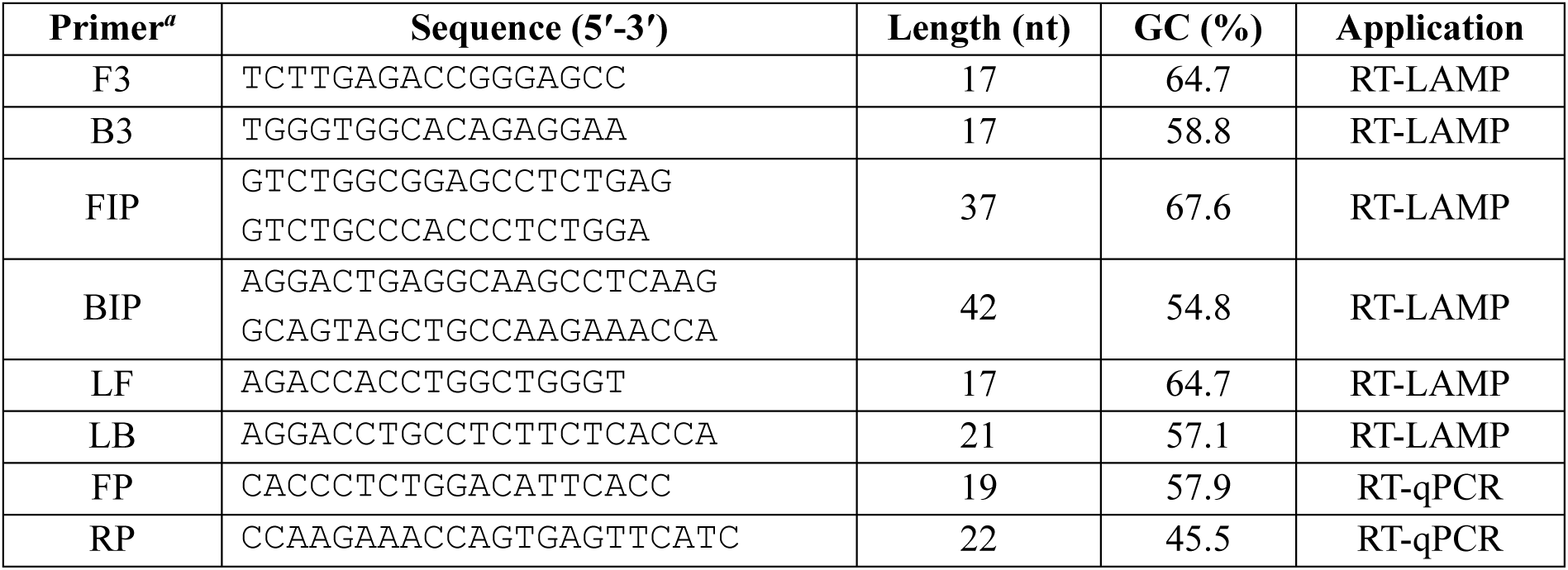
*KISS1* mRNA-specific primer sequences used for RT-LAMP and qPCR assays.

### 2.9 Complementary DNA (cDNA) preparation and RT-qPCR amplification conditions

Following extraction of exosomal total RNA, complementary DNA (cDNA) was synthesized using the QuantiTect Reverse Transcription Kit (Qiagen) according to the manufacturer’s protocol. Briefly, RNA samples were first treated with genomic DNA (gDNA) Wipeout Buffer to remove genomic DNA. The RNA was then mixed with QuantiTect Reverse Transcription Master Mix containing reverse transcriptase, RT buffer, and RT primer mix, and the reaction volume was adjusted to 20 µL with nuclease-free water. Reverse transcription was performed in a CFX Opus 96 Real-Time PCR System (Bio-Rad Laboratories, USA) at 42 °C for 15 min, followed by 95 °C for 3 min to inactivate the enzyme. The resulting cDNA was diluted with nuclease-free water and stored at −20 °C until further analysis.

Using the synthesized cDNA as the template, quantitative PCR (qPCR) was performed on the CFX Opus 96 Real-Time PCR System. The amplification protocol consisted of an initial denaturation step at 95 °C for 2 min, followed by 40 amplification cycles. Each cycle included denaturation at 95 °C for 5 s, annealing at 61 °C for 10 s, and extension at 72 °C for 20 s, with fluorescence signals recorded at the end of each extension step (plate read). After amplification, melting curve analysis was conducted to confirm product specificity by gradually increasing the temperature from 60 °C to 95 °C with a ramp rate of 0.5 °C/s. All reactions were performed in technical triplicates.

### 2.10 RT-LAMP reaction conditions

RT-LAMP of KISS1 mRNA was performed using WarmStart Colorimetric LAMP 2X Master Mix (DNA & RNA) (NEB, England) in a total reaction volume of 25 µL. Each reaction mixture contained 12.5 µL of the 2X master mix, 2.5 µL of 10X LAMP primer mix (final concentrations: 2 µM F3 and B3, 16 µM FIP and BIP, and 4 µM LF and LB), 0.5 µL SYBR Green I dye, 2-9.5 µL of mRNA template, and a proper amount of nuclease-free water to adjust the final volume to 25 µL. After all reagents and target templates were added at the appropriate volumes, amplification was carried out at 65 °C for 40-60 min in a preheated CFX Opus 96 Real-Time PCR System (Bio-Rad Laboratories, USA), and fluorescence signals were monitored in real time.

### 2.11 Agarose gel electrophoresis

A 1.5% agarose gel was prepared in 1X TAE buffer, with 10 µL SYBR Safe DNA stain (Invitrogen, Australia) added per 100 mL of gel solution. For electrophoresis, 5 µL of sample was mixed with 1 µL of 6X gel loading dye (Thermo Fisher Scientific, USA) and loaded onto the gel. Electrophoresis was performed at 90 V for 60 min. After separation, DNA bands were visualized and imaged using the Bio-Rad GelDoc Go Imaging System.

### 2.12 Statistical analysis

All statistical analyses and graphical plots were performed using GraphPad Prism v10 (GraphPad Software, USA). Threshold times and quantification cycle (Cq) values were expressed as mean ± standard deviation (SD) from three independent replicates. Differences between groups were assessed using an unpaired nonparametric Mann-Whitney U test, and statistical significance was defined as *p* < 0.05. Simple linear regression analysis was used to generate standard curves by plotting the threshold time for RT-LAMP (or Cq value for RT-qPCR) against the logarithm of input mRNA copies per reaction. For comparisons among more than two independent groups, data were analyzed in lognormal one-way Brown-Forsythe and Welch ANOVA, followed by Dunnett’s T3 multiple-comparisons test with adjusted *p* values for all pairwise comparisons. The limit of detection (LOD) was calculated using the following equation (1) [30, 31]:

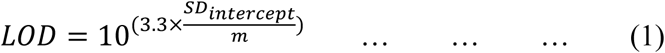

where, *m* is the slope of regression equation and *SD_intercept_* represents the standard deviation of the intercept which was calculated from another equation as expressed in equation (2).

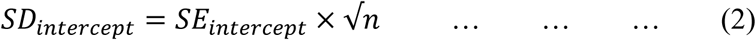

where, *SE_intercept_* is the standard error of the intercept and *n* is the total number of reactions included in the regression analysis. The limit of quantification (LOQ) was defined as the lowest concentration that could be quantified with acceptable precision, expressed as a coefficient of variation (CV) of ≤25% [32]. CV was calculated using the equation (3) from replicate measurements.

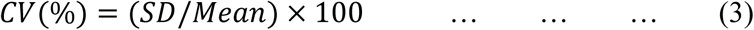

## 3. RESULTS AND DISCUSSION

### 3.1 Assay principle

Figure 1 illustrates the overall Exo-LAMP assay principle for selective, extraction-free detection of placental exosomal KISS1 mRNA. (i-ii) The assay starts with exosome-depleted serum spiked PLAP^P^ EVs or conditioned culture medium (CCM) containing PLAP^P^ EVs, which are selectively captured using anti-PLAP antibody-conjugated magnetic beads within 20 minutes. This exosome isolation step is simpler and faster than conventional kit-based bulk exosome isolation, which typically requires about 14 h. (iii) A magnet is then used to separate the exosome-bound anti-PLAP antibody conjugated magnetic beads from the surrounding medium, and the supernatant is removed. (iv) The captured exosomes are resuspended in RNase inhibitor containing phosphate-buffered saline (PBS) and lysed at 90 °C for 5 minutes, allowing direct release of KISS1 mRNA and other nucleic acids without RNA extraction kit. (v-vii) The colorimetric RT-LAMP reagent and KISS1-specific primer mix are then added directly to the same tube. The reaction proceeds for 25-30 minutes as a one-pot isothermal amplification assay for semi-quantitative detection of KISS1 mRNA. As RT-LAMP is performed at a constant temperature, the workflow avoids conventional thermocycler-based RT-qPCR, which usually takes about 2 h. Positive amplification is detected in real time and by a visible color change from pink to yellow. Overall, Exo-LAMP provides a simple one-tube, extraction-free, and rapid workflow for exosomal mRNA detection within 50 – 55 minutes, compared with the conventional kit-based workflow that requires about 16 h.

**Figure 1.**
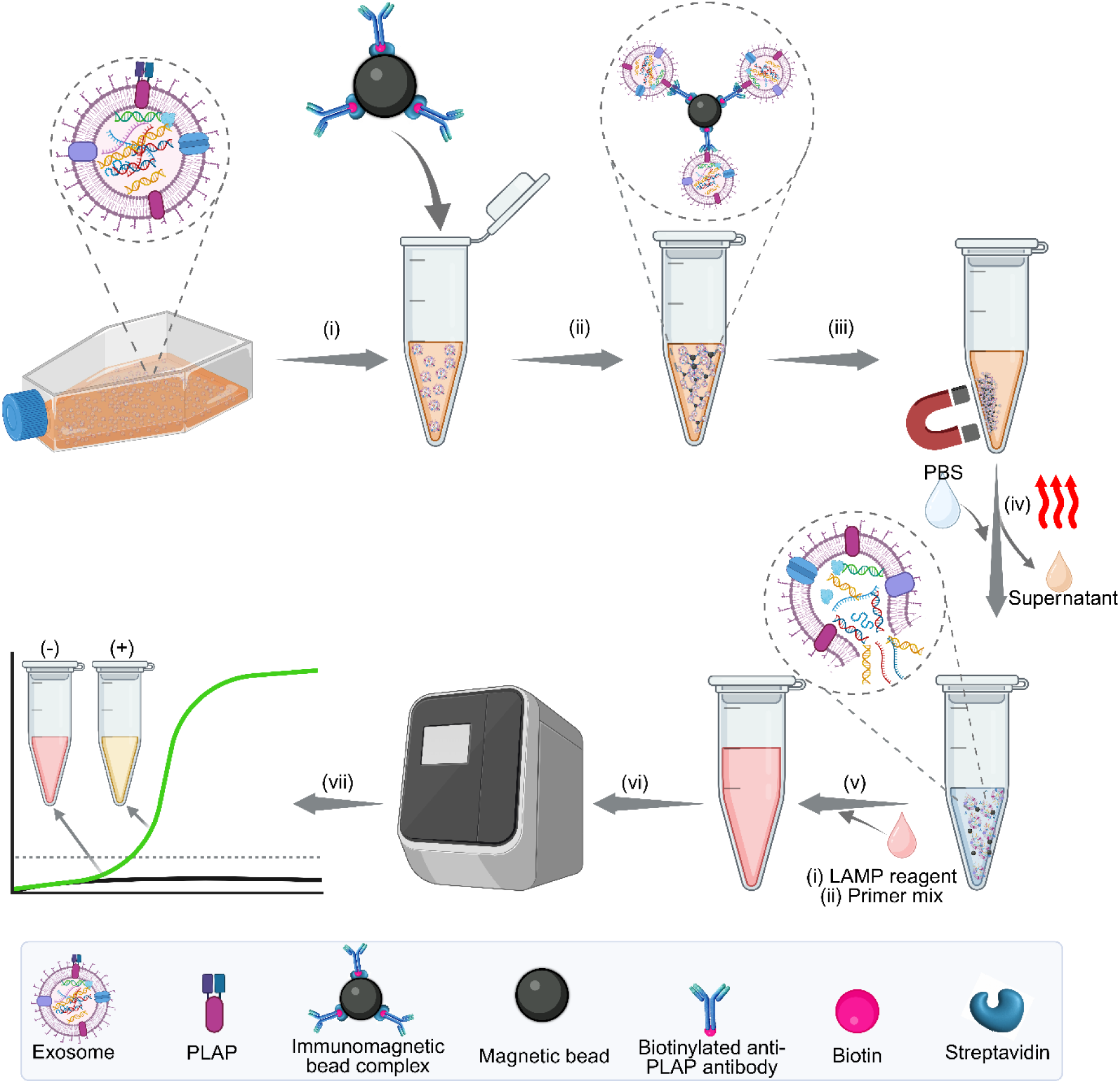
Schematic workflow of the Exo-LAMP assay platform. (i–ii) Placental alkaline phosphatase positive (PLAPP) exosomes in exosome-depleted serum-spiked samples or conditioned culture medium (CCM) are captured using anti-PLAP antibody-conjugated magnetic beads within 20 minutes. (iii–iv) The antibody–bead–exosome complexes are separated using a magnet, resuspended in RNase inhibitor containing PBS, and heated at 90 °C for 5 minutes to release KISS1 mRNA without the need for RNA extraction kits. (v–vi) RT- LAMP reagents and KISS1-specific primers are added directly to the same tube for one-pot amplification within 25–30 minutes. (vii) Positive amplification is detected in real time via amplification curves and a visible color change from pink to yellow.

### 3.2 Exosome characterization and immunomagnetic capture efficiency of anti-PLAP

To characterize EVs isolated from the PLAP^P^ BeWo and PLAP^N^ HTR-8 cell lines, nanoparticle tracking analysis (NTA) was performed. As shown in Figures 2A and 2B, EVs from both cell lines showed nanoscale size distributions within the expected exosomal range of 30-150 nm [10]. BeWo-derived vesicles showed a dominant peak at around 90 nm, with a mean particle size of 90.25 ± 4.23 nm, a mode of 85.19 nm, and a concentration of 16.90 ± 1.54 billion particles/mL. Similarly, HTR-8-derived vesicles showed a comparable distribution, with a mean diameter of 98.40 ± 4.89 nm, a mode of 93.35 nm, and a concentration of 12.40 ± 1.02 billion particles/mL. To further assess the physicochemical properties of the isolated exosomes, zeta potential analysis was performed (Figure 2C). Isolated bare exosomes showed a moderately negative surface charge (−13.2 ± 0.63 mV), which is consistent with the anionic phospholipid composition of EV membranes. The observed zeta potential also falls within the range commonly reported for exosomes (−10.42 to −15.84 mV), supporting the presence of vesicles with typical surface characteristics. This finding further indicates that the progenitor cells from which these vesicles originated retained relatively intact membrane properties without obvious pathological disruption [33].

**Figure 2.**
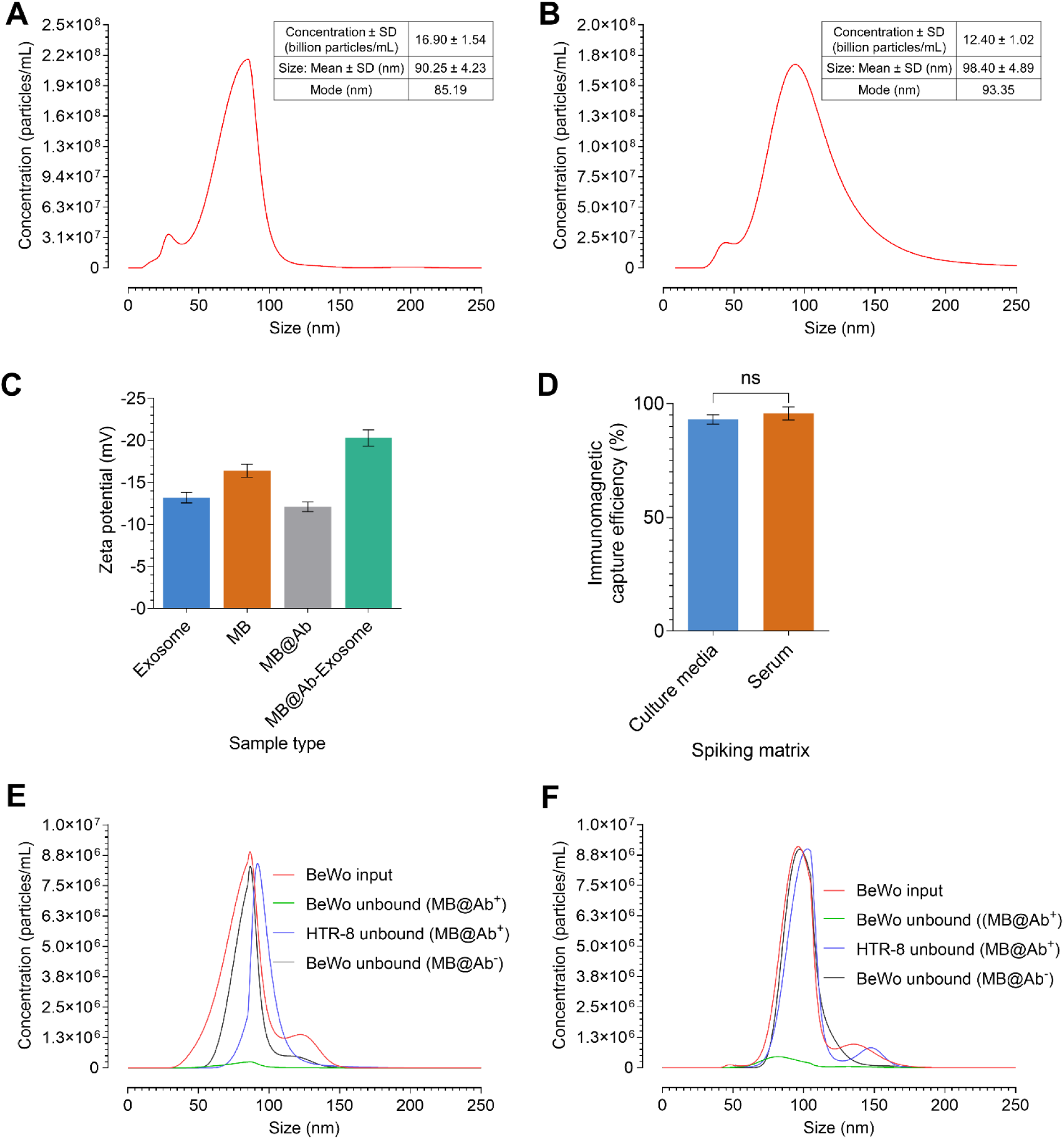
Characterization of extracellular vesicles (EVs) and evaluation of immunomagnetic capture. (A–B) Nanoparticle tracking analysis (NTA) size distribution profiles of EVs derived from PLAP^P^ BeWo cells (A) and PLAP^N^ HTR-8 cells (B), showing a typical exosomal size range of 30–150 nm. (C) Zeta potential analysis confirming antibody (Ab) immobilization on magnetic beads (MB) and formation of MB@Ab–exosome complexes. (D) Comparison of capture efficiency of MB@Ab for BeWo-derived exosomes in CCM and exosome-depleted serum, showing comparable immunomagnetic enrichment with no significant difference between the two matrices (p = 0.40). (E–F) Capture efficiency of MB@Ab evaluated by NTA after spiking BeWo-derived exosomes into culture medium (E, 95.66 ± 2.89%) and exosome-depleted serum (F, 93.04 ± 2.10%), demonstrating efficient immunomagnetic enrichment across different sample matrices. [Terms used in the figure: BeWo input, initial BeWo-derived exosome population prior to immunomagnetic capture; BeWo unbound (MB@Ab^+^), BeWo-derived exosomes remaining in the supernatant after capture with anti-PLAP antibody-functionalized magnetic beads; HTR-8 unbound (MB@Ab^+^), HTR-8- derived exosomes remaining in the supernatant after capture with anti-PLAP antibody- functionalized magnetic beads; BeWo unbound (MB@Ab^−^), BeWo-derived exosomes remaining in the supernatant without antibody-functionalized magnetic beads (negative control).]

To define the biological identity of these exosomes, Western blot analysis was performed using PLAP as a marker (Figure S1). PLAP was detected only in BeWo-derived exosomes, while no PLAP signal was observed in HTR-8-derived exosomes. This finding agrees with the known PLAP^P^ phenotype of syncytiotrophoblast-derived BeWo cells and the PLAP^N^ status of extravillous trophoblast HTR-8 cells [20, 34]. Together, the NTA, zeta potential, and Western blot results show that although both cell lines released vesicles with similar exosome-like size profiles, only BeWo-derived exosomes carried the placental marker PLAP. This supports the use of BeWo-derived exosomes as a suitable model for PLAP-targeted placental exosome isolation and subsequent KISS1 mRNA analysis.

To confirm immobilization of the anti-PLAP antibody on magnetic bead (MB) and the subsequent binding of exosomes to the anti-PLAP antibody-functionalized beads (MB@Ab), zeta potential measurements were performed (Figure 2C). While the isolated bare exosomes showed a moderately negative surface charge (−13.2 ± 0.63 mV), the streptavidin-coated MB exhibited a slightly more negative zeta potential (−16.4 ± 0.79 mV). After immobilization of the biotinylated anti-PLAP antibody on the MB surface, the zeta potential of MB@Ab shifted to −12.1 ± 0.58 mV, indicating successful surface functionalization [35]. Following incubation with exosomes, the MB@Ab-exosome complexes showed a further shift toward a more negative value (−20.3 ± 0.97 mV), consistent with the binding of negatively charged vesicle membranes to the MB@Ab [36].

To further evaluate the exosome capture efficiency of MB@Ab, a known number of exosomes was spiked into fresh culture medium, and exosome-depleted serum (Figures 2D-F). After incubation with MB@Ab, the remaining vesicles in the supernatant were quantified by NTA to determine the proportion of unbound particles. In both matrices, only a small fraction [Figure 2E, green curve, BeWo unbound (MB@Ab^+^); Figure 2F, green curve, BeWo unbound (MB@Ab^+^)] of the initial BeWo-derived exosomal input [Figure 2E, red curve, BeWo input; Figure 2F, red curve, BeWo input] remained in the supernatant after incubation with MB@Ab. This finding indicates efficient immunomagnetic capture of PLAP^P^ BeWo-derived exosomes by the anti-PLAP antibody-functionalized beads. The capture efficiency is found 95.66 ±2.89% in culture medium and 93.04 ± 2.10% in serum, which is comparable to other PLAP-targeted immunoaffinity-based exosome capture systems [20, 34]. In contrast, when BeWo-derived exosomes were incubated with non-functionalized beads lacking antibody (MB@Ab^−^), the supernatant profiles [Figure 2E, black curve, BeWo unbound (MB@Ab^−^); Figure 2F, black curve, BeWo unbound (MB@Ab^−^)] remained largely similar to the initial BeWo input, indicating very minimal nonspecific capture in the absence of anti-PLAP antibody.

Likewise, PLAP^N^ HTR-8-derived exosomes showed little reduction in the unbound fraction after incubation with MB@Ab^+^, and the supernatant profile [Figure 2E, blue curve, HTR-8 unbound (MB@Ab^+^); Figure 2F, blue curve, HTR-8 unbound (MB@Ab^+^)] remained similar to the input [Figure 2E, red curve, BeWo input; Figure 2F, red curve, BeWo input], with only a slight shift toward larger particle size, consistent with the size distribution observed for HTR-8-derived exosomes in Figure 2B. This suggests that the MB@Ab^+^ was highly selective for PLAP^P^ exosomes and showed no significant interaction with PLAP^N^ vesicles. Importantly, capture performance was found to be almost similar in both fresh culture medium and exosome-depleted serum. The comparable efficiencies across these two matrices (95.66 ± 2.89% in CCM and 93.04 ± 2.10% in exosome-depleted serum) indicate that the anti-PLAP- functionalized MB@Ab system maintained selective binding even in the more complex serum environment (Figure 2D). These findings support the robustness of the immunomagnetic platform and its potential applicability to clinically relevant samples, where target exosome isolation is often hindered by competing biomolecules and non-target bulk vesicles.

### 3.3 Optimization of RT-LAMP reaction conditions

#### 3.3.1. Reaction temperature

To identify the optimal reaction temperature for the RT-LAMP, temperature gradient experiments were performed. *Bacillus stearothermophilus* polymerase 2.0 (*Bst* 2.0 WarmStart, NEB #M0538) is reported to perform optimally within the range of 60-70 °C. However, as amplification efficiency decreases at the higher end of this range (68-70 °C) [37], the reaction was evaluated across a gradient from 60.0 °C to 66.0 °C. Real-time fluorescence monitoring showed that amplification occurred between 62.4 °C and 66.0 °C (Figure S2A). When the RT-LAMP amplicons were further analyzed by agarose gel electrophoresis, the characteristic LAMP bands were observed for reactions performed at 62.4 °C to 66 °C, whereas no bands were detected at 60 °C to 61.2 °C or in the no-template control (Figure S2B). Based on the combined fluorescence and gel electrophoresis results, 65 °C was selected as the optimal reaction temperature and was used for all subsequent RT-LAMP experiments.

#### 3.3.2. Primer specificity for KISS1 mRNA

As exosomes can contain genomic DNA (gDNA) in addition to RNA, it is important to confirm that the signal obtained in this semi-quantitative mRNA reaction does not arise from contaminating KISS1 genomic sequence. To address this, the LAMP primers were designed to span exon-exon junctions [38], thereby preventing amplification from the genomic KISS1 DNA sequence and ensuring selective detection of the mRNA target.

Primer specificity was evaluated using synthetic KISS1 mRNA tested alone and in combination with gDNA isolated from the BeWo cell line. As shown by the real-time amplification profiles (Figure S3A), strong amplification was observed in reactions containing KISS1 mRNA alone and in reactions containing KISS1 mRNA mixed with gDNA, whereas no amplification was detected in reactions containing only gDNA or in the NTC. No amplification from the NTC supports that the reaction was free from detectable contamination and carryover amplification. Carryover contamination was prevented by using the WarmStart Colorimetric LAMP 2X Master Mix with UDG (NEB #M1804), which contains deoxyuridine triphosphate (dUTP) and Antarctic Thermolabile Uracil-DNA Glycosylase (UDG) to prevent amplification of contaminating products carried over from previous reactions (https://www.neb.com/en-au/products/m1804-warmstart-colorimetric-lamp-2x-master-mix-with-udg). In this system, any carryover amplicons from earlier reactions that contain deoxyuridine are degraded by UDG before amplification starts. The enzyme is then inactivated during the 65 °C reaction, so newly formed products are not affected.

The specificity of the amplified products was further supported by melt curve and melt peak analyses. Reactions containing KISS1 mRNA alone and KISS1 mRNA plus gDNA showed highly similar melt profiles (Figure S3B) and a single dominant melt peak (Figure S3C), consistent with amplification of the same specific target product under both conditions. The colorimetric end-point results were fully consistent with the fluorescence-based data (Figure S3D), with positive reactions changing from pink to yellow in the KISS1 mRNA (Tube 1) and KISS1 mRNA plus gDNA (Tube 3) reactions, while the gDNA-only (Tube 2) and NTC reactions remained pink (Tube 4).

These findings confirm that the RT-LMAP reaction remained specific for KISS1 mRNA, with no detectable amplification from KISS1 genomic sequences even in the presence of a complex gDNA background, and no evidence of nonspecific amplification due to contamination or carryover.

### 3.4 Effect of sample matrix on KISS1 mRNA detection

To assess the analytical performance of the RT-LAMP reaction under biologically relevant conditions, synthetic KISS1 mRNA transcripts were serially diluted and spiked into three different matrices: tris-EDTA (TE) buffer, exosome-depleted serum, and RNase inhibitor-treated exosome-depleted serum.

When synthetic KISS1 mRNA was spiked into TE buffer, the RT-LAMP produced rapid amplification and clearly distinguishable signals across serial dilutions ranging from 10^1^ to 10^8^ copies per reaction (Figure 3A). The standard curve showed a strong linear relationship (coefficient of correlation, *r^2^* = 0.9891) between threshold time (Tt) and the log_10_-transformed mRNA copy number (Figure 3B). Based on this regression, the theoretical limit of detection (LOD) was estimated to be approximately 1 copy per reaction, consistent with the high analytical sensitivity observed in our previous LAMP-based studies [30]. The lowest concentration showing a coefficient of variation (CV) of ≤25% was defined as the limit of quantification (LOQ) [32], which was determined to be 10 copies per reaction.

**Figure 3.**
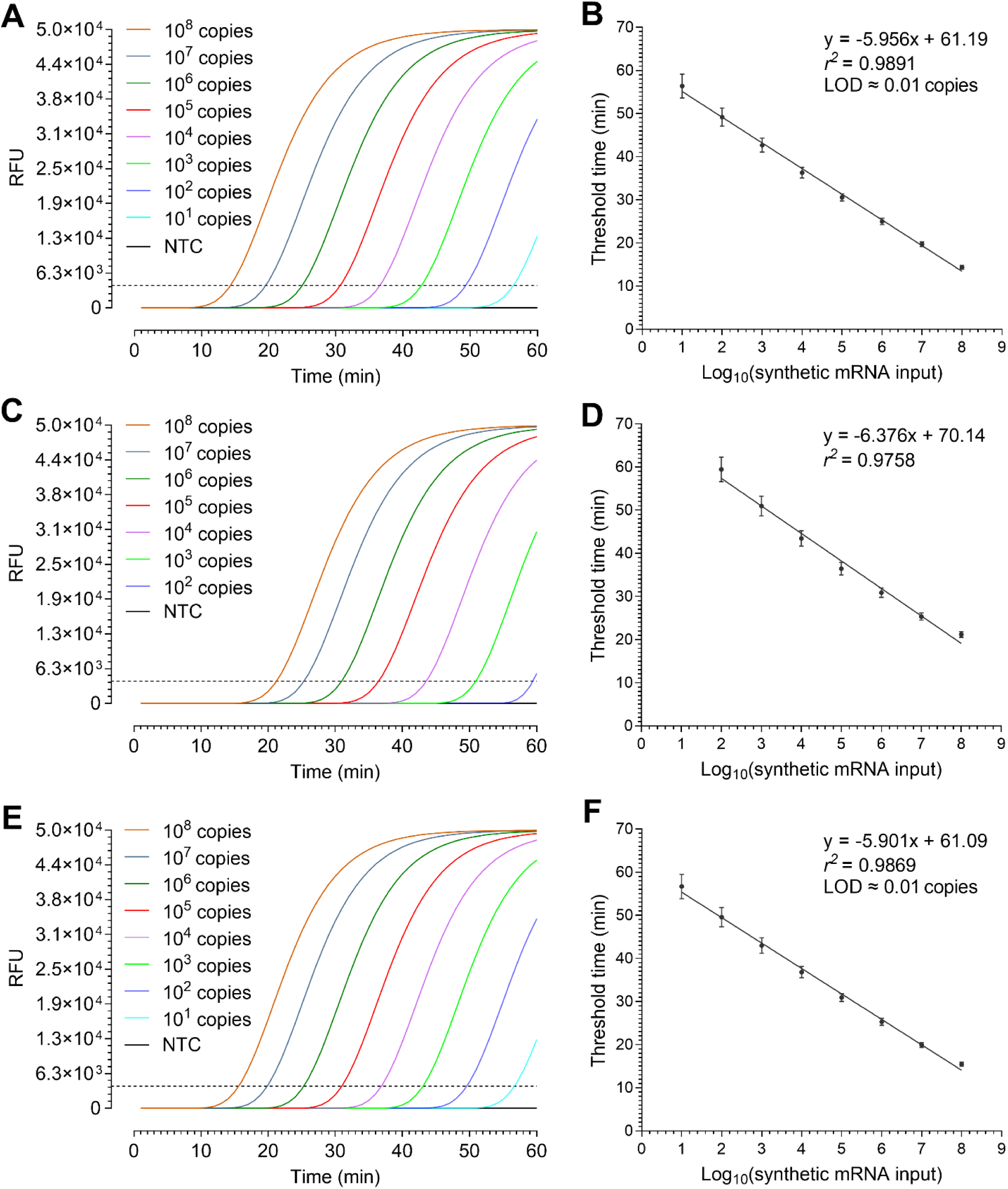
Analytical performance of RT-LAMP for KISS1 mRNA detection across different sample matrices. (A) Real time RT-LAMP amplification curves obtained from serial dilutions of synthetic KISS1 mRNA (10^1^ to 10^8^ copies per reaction) spiked into TE buffer. (B) Corresponding calibration curve showing the relationship between threshold time and log_10_- transformed mRNA copy number. (C) Amplification curves for synthetic KISS1 mRNA spiked into exosome-depleted serum under identical RT-LAMP conditions. (D) Corresponding calibration curve for panel C. (E) Amplification curves for synthetic KISS1 mRNA spiked into exosome-depleted serum pretreated with RNase inhibitor. (F) Corresponding calibration curve for panel E.

To evaluate the effect of the protein- and enzyme-rich serum environment on reaction performance, synthetic KISS1 mRNA was spiked into exosome-depleted human serum and analyzed under the same RT-LAMP conditions. The RT-LAMP reaction still showed consistent amplification behavior across the dilution series (Figure 3C), indicating that the target remained detectable in this more complex biological matrix. However, compared with reactions performed in TE buffer (Figure 3A), the Tts were slightly delayed, and no amplification was detected at input levels below 10^2^ copies per reaction (Figure 3C-D). This reduction in sensitivity suggests a matrix-dependent inhibitory effect, most likely due to serum-associated components that interfere with amplification efficiency, including endogenous RNases [8]. Because serum is known to contain high levels of RNase activity [39], partial degradation of the target RNA is likely to have contributed to the reduced reaction sensitivity observed in this matrix [8].

To examine if RNA degradation in serum contributed to reduced reaction performance, serum samples were first treated with RiboLock RNase Inhibitor 0.1 U/µL (Thermo Scientific, #EO0381) before spiking with KISS1 mRNA. Under this condition, the amplification curves showed faster and more consistent reaction kinetics (Figure 3E) than those observed in untreated serum samples (Figure 3C), indicating that inhibition of RNase activity improved target stability and reaction performance [26]. Notably, the RT-LAMP detected as few as 10 copies of mRNA per reaction, restoring the dynamic range to 10^1^-10^8^ copies per reaction, which was comparable to that observed when the same mRNA targets were spiked into TE (Figure 3A). The standard curve showed a strong linear relationship between threshold time and the log_10_-transformed mRNA copy number (*r^2^* = 0.9869) (Figure 3F). The LOD was recovered to approximately 1 copy per reaction, and the LOQ was again determined to be 10 copies per reaction, matching the values obtained in the case of TE buffer matrix (Figure 3B).

### 3.5 Heat-mediated lysis of isolated exosomes

To optimize heat-mediated lysis for direct downstream analysis of exosomal nucleic acid, a defined amount of BeWo cell-derived exosomes (1.7 × 10^6^ particles) was suspended in exosome-depleted serum pretreated with RNase inhibitor and exposed to four thermal conditions: 90 °C for 5 minutes, 90 °C for 10 minutes, 95 °C for 5 minutes, and 95 °C for 10 minutes [23, 24]. The resulting lysates were used directly for RT-LAMP under optimized conditions. KISS1 mRNA copy number was then semi-quantitatively estimated from the Tt using the equation (y = −5.901x + 61.09) shown in Figure 3F and compared with the yield obtained using a commercial RNA extraction kit. As shown in Figure 4A, all heat-lysis conditions generated broadly comparable amplification profiles, although the 90 °C for 5 minutes condition most closely resembled the amplification pattern obtained with kit-extracted RNA. This trend was also reflected in the Tt results (Figure 4B), where only modest variation was observed across conditions. While the differences in amplification behavior and Tt were not pronounced, the estimated KISS1 mRNA copy number revealed a clearer distinction between lysis conditions (Figure 4C). Among the tested conditions, heat lysis at 90 °C for 5 minutes produced the highest copy number and was not significantly different from the kit-based extraction, whereas the other heating conditions yielded significantly lower mRNA copy number.

**Figure 4.**
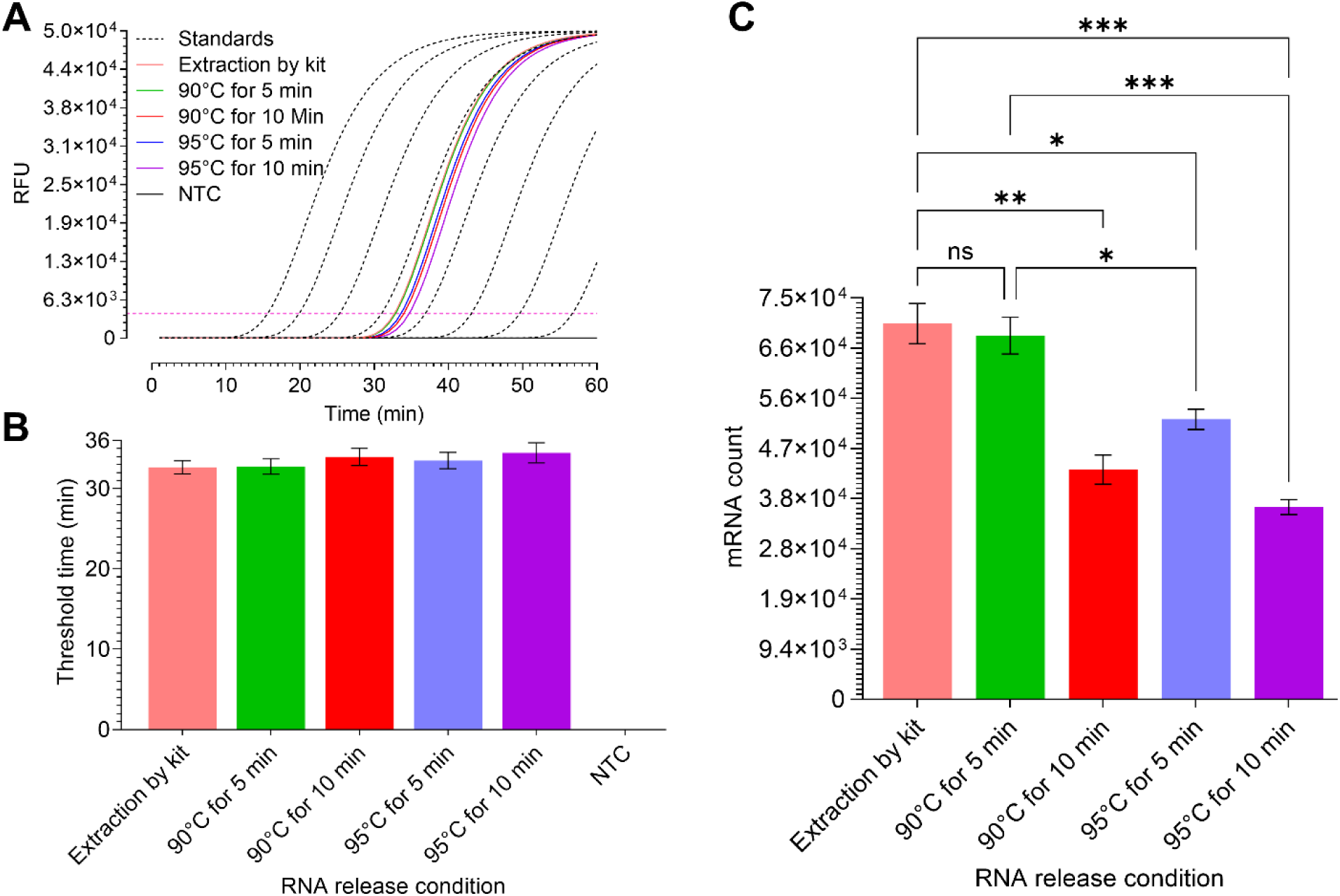
Optimization of heat-mediated lysis for exosomal KISS1 mRNA analysis. (A) RT- LAMP amplification curves generated from BeWo cell-derived exosomes after heat lysis under different temperature and time conditions, with the extraction kit-based control included for comparison. (B) Threshold time (Tt) values for the corresponding lysis conditions and the extraction kit-based control. (C) Semi-quantitative KISS1 mRNA copy number estimated from Tt values using the standard curve shown in Figure 3F. Overall, heat lysis at 90 °C for 5 minutes showed the closest performance to kit-based extraction.

A similar direct heat-lysis strategy has been reported previously, where enriched exosomes were heated at 95 °C for 10 minutes and the released RNA was used directly for downstream isothermal analysis without further purification [24]. Another study optimizing EV miRNA extraction identified 90 °C for 5 minutes as the optimal lysis temperature [23]. Consistent with the latter study [23], our results showed that lysis at 90 °C for 5 minutes gave the best overall performance and was therefore selected as the optimal condition for subsequent analysis. This heat treatment also reduces nuclease activity [25], but released RNA can still remain vulnerable to RNase-mediated degradation during the heating transition [8], therefore, RNase inhibitors added into serum help preserve RNA integrity and improve downstream detection reliability [26]. Traditional nucleic acid extraction methods, including silica column-based kits, are relatively time-consuming, typically requiring about 1.5 h, multiple processing steps, and an estimated per-test cost of approximately USD 12 [40]. By contrast, this one-step heat-mediated lysis method requires only 5 minutes and eliminates the need for a commercial extraction kit, highlighting its potential as a rapid, simple, and low-cost alternative for nucleic acid extraction.

When immediate processing was not feasible, isolated exosomes were stored under controlled conditions. In accordance with the Minimal Information for Studies of Extracellular Vesicles 2023 (MISEV2023) guidelines [41], repeated freeze–thaw cycles were minimized to preserve vesicle integrity. When storage was required within the workflow, exosomes were maintained at −80 °C in conditioned culture medium supplemented with dimethyl sulfoxide (DMSO) and were subjected to no more than two freeze–thaw cycles. The effects of storage conditions on exosome stability are presented in Figure S4 of the Supporting Information.

### 3.6 Analytical performance of Exo-LAMP assay

After optimizing all of the above-described steps of the Exo-LAMP platform, its overall performance was evaluated using exosomes spiked into exosome-depleted serum. In the Exo- LAMP assay, serial dilutions containing defined numbers of exosomes (10^3^ to 10^8^ per reaction) were captured using the anti-PLAP antibody-containing MB@Ab system, followed by lysis at 90 °C for 5 minutes to release KISS1 mRNA, and the released KISS1 mRNA was then analyzed by RT-LAMP. The same dilution series was also analyzed using the conventional workflow, which included kit-based exosome isolation and RNA extraction, followed by RT-qPCR based mRNA semi-quantification.

The Exo-LAMP assay produced clear amplification across dynamic range of exosome inputs from 10^3^ to 10^8^ (Figure 5A). No amplification was detected from PLAP^N^ HTR-8-derived exosomes, confirming the assay specificity for the targeted PLAP^P^ exosome population. The colorimetric results showed the same detection range (Figure 5B), with positive reactions observed from only 10^3^ to 10^8^ exosomes. This range is consistent with exosome concentrations commonly found in biological fluids (1 × 10^5^ to 3 × 10^9^ exosomes/µL) [42], which supports the practical use of the Exo-LAMP assay for exosomal mRNA analysis. By comparison, the conventional RT-qPCR workflow showed a narrower detection range, from 10^4^ to 10^8^ exosomes, with no reliable signal below 10^4^ exosomes per reaction. This lower sensitivity was also reflected in the estimated mRNA counts (Figure 5C). Using the assay-specific standard curves and regression equations developed for each platform (Figures 3E-F and y = −5.901x + 61.09 for Exo-LAMP; Figures S5A-C and y = −4.290x + 45.68 for conventional RT-qPCR), KISS1 mRNA abundance was estimated for each exosome input.

**Figure 5.**
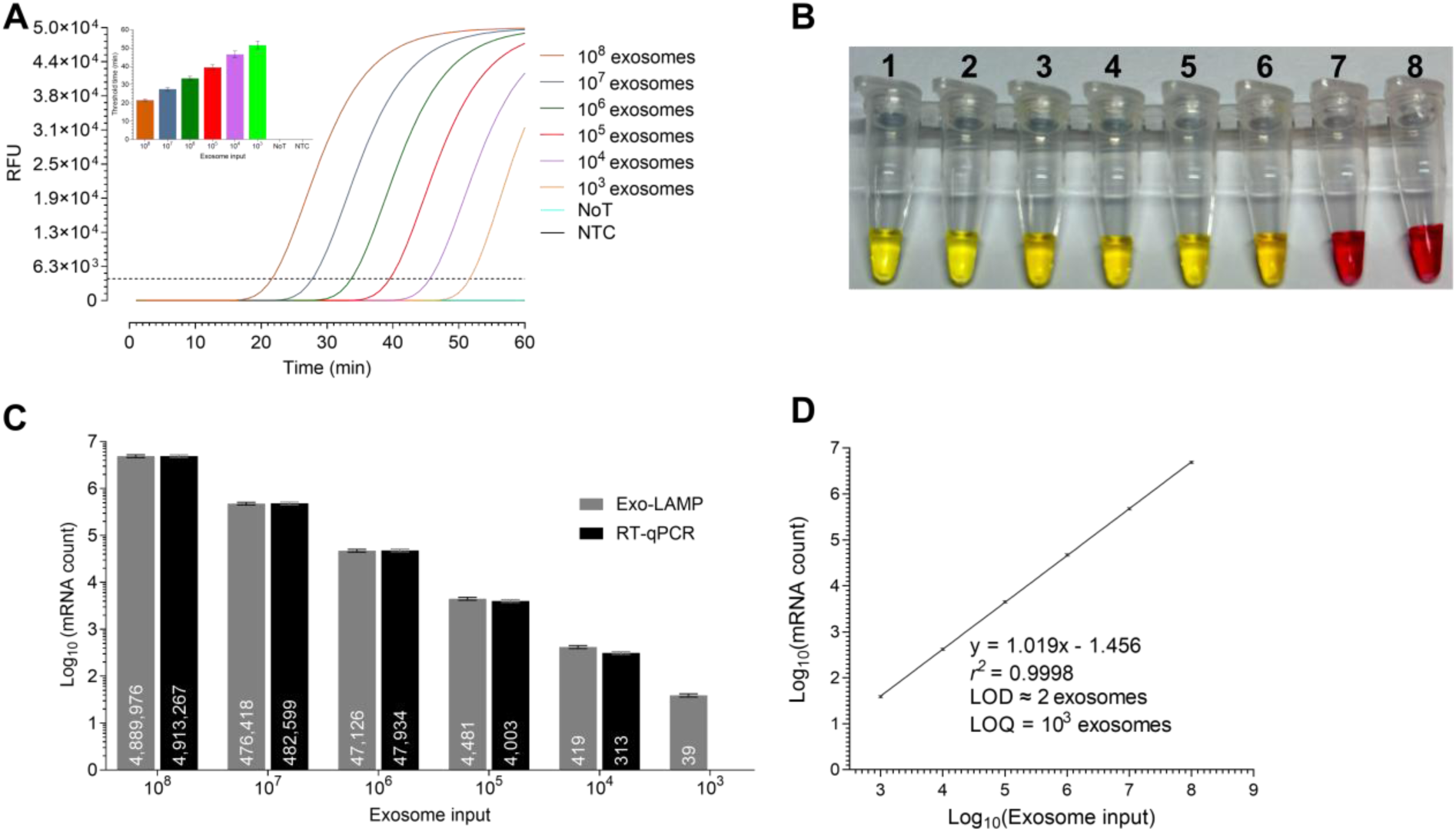
Analytical response of Exo-LAMP for KISS1 mRNA quantification as a function of exosome input. (A) Real time amplification curves for exosome inputs ranging from 10^3^ to 10^8^ per reaction. (B) Corresponding colorimetric readout. (C) Estimated KISS1 mRNA copy numbers obtained by Exo-LAMP and conventional RT-qPCR across the exosome dilution series; values within bars indicate calculated copy numbers. (D) Linear correlation between estimated KISS1 mRNA copy number and exosome input for Exo-LAMP, with a limit of detection of approximately 2 exosomes and a limit of quantification of 10^3^ exosomes.

According to Figure 5C, Exo-LAMP and conventional RT-qPCR produced comparable mRNA counts across the higher input range (10^6^ to 10^8^ exosomes), with RT-qPCR even giving slightly higher values. However, as exosome input decreased (10^5^ to 10^3^), the RT-qPCR-derived mRNA counts declined more sharply than those obtained by Exo-LAMP, and no mRNA count could be detected by RT-qPCR at 10^3^ exosome input, whereas Exo-LAMP still produced a measurable signal. This pattern suggests that the integrated Exo-LAMP assay retained stronger sensitivity when the target became scarce. One likely reason is the difference in workflow structure. The conventional method requires several separate steps, including exosome isolation, RNA extraction, transferring for reverse transcription, and downstream amplification. When the starting exosome number is low, each additional handling step may lead to partial sample loss and reduced RNA recovery [22, 23]. These losses may have only a small effect at high exosome input, but they become much more important as the available target exosomes decrease. In contrast, the Exo-LAMP workflow reduces sample manipulation by combining direct in-tube RNA release and reverse transcription integrated isothermal amplification into a more continuous process. This simpler workflow likely helps preserve low- abundance RNA and reduces the opportunity for material loss before amplification [22].

Furthermore, both Exo-LAMP and the conventional RT-qPCR-based workflow demonstrated mRNA counting capability, with estimated LODs of approximately 2 and 4 exosomes, respectively (Figures 5D and S5D), whereas Exo-LAMP achieved a lower LOQ of 10^3^ exosomes compared with 10^4^ exosomes for RT-qPCR. These results indicate that although both workflows were similarly capable of detecting very low target levels, Exo-LAMP maintained reliable mRNA semi-quantification across a more lower exosome input range than that of conventional RT-qPCR-based method. This interpretation was further supported by the stoichiometric semi-quantitative ratio of mRNA and exosomes, performed using the assay- specific regression equations [y = 1.019x − 1.456 (Figure 5D) for Exo-LAMP; y = 1.047 x − 1.652 (Figure S5D) for RT-qPCR]. Based on these calculations, Exo-LAMP yielded an estimated ratio of approximately 1 KISS1 mRNA molecule per 27 exosomes, whereas the conventional RT-qPCR workflow yielded approximately 1 mRNA molecule per 38 exosomes. Previous RNA-sequencing analyses have suggested that EV-associated mRNAs are often fragmented and that intact transcripts are generally present at very low abundance [43]. In glioma stem cell-derived EV preparations, stoichiometric analysis estimated approximately one mRNA or mRNA fragment per 10 EVs on average, whereas individual mRNA species were much rarer. The TMSB10 mRNA was detected at approximately one copy per 1000 EVs, while ACTB and GAPDH were present at about one copy per 10,000 EVs. Disease-relevant glioma-associated mRNAs were even less abundant, with COL1A2, IDH1, and EGFR detected at approximately one copy per 100,000 EVs and TP53 and PTEN at approximately one copy per 1,000,000 EVs [43].

### 3.7. Evaluating the performance of Exo-LAMP in the presence of non-target exosomes

After establishing the dynamic detection range of the Exo-LAMP assay across exosome inputs from 10^3^ to 10^8^, the assay was further assessed for its ability to quantify low-abundance PLAP^P^ exosomes within heterogeneous mixtures containing excess non-target exosomes. For this purpose, exosome-depleted serum-spiked exosome mixtures were prepared by combining PLAP^P^ BeWo-derived exosomes with PLAP^N^ HTR-8-derived exosomes at target-to-non-target ratios of 100:0, 50:50, 25:75, 10:90, and 1:99. These mixtures corresponded to BeWo-derived exosome inputs of 1.7 × 10^5^, 8.5 × 10^4^, 4.25 × 10^4^, 1.7 × 10^4^, and 1.7 × 10^3^, respectively. The resulting KISS1 mRNA signals were compared with the expected mRNA counts predicted from the Exo-LAMP regression model (y = 1.019x – 1.456; Figure 5D), showing that the experimentally measured counts remained close to the predicted values across all mixing ratios (Figure 6A). This agreement was maintained even at the 1:99 condition. Although the measured value (48) at this lowest ratio was lower than the expected value (54), the assay still produced a clear and quantifiable signal.

**Figure 6.**
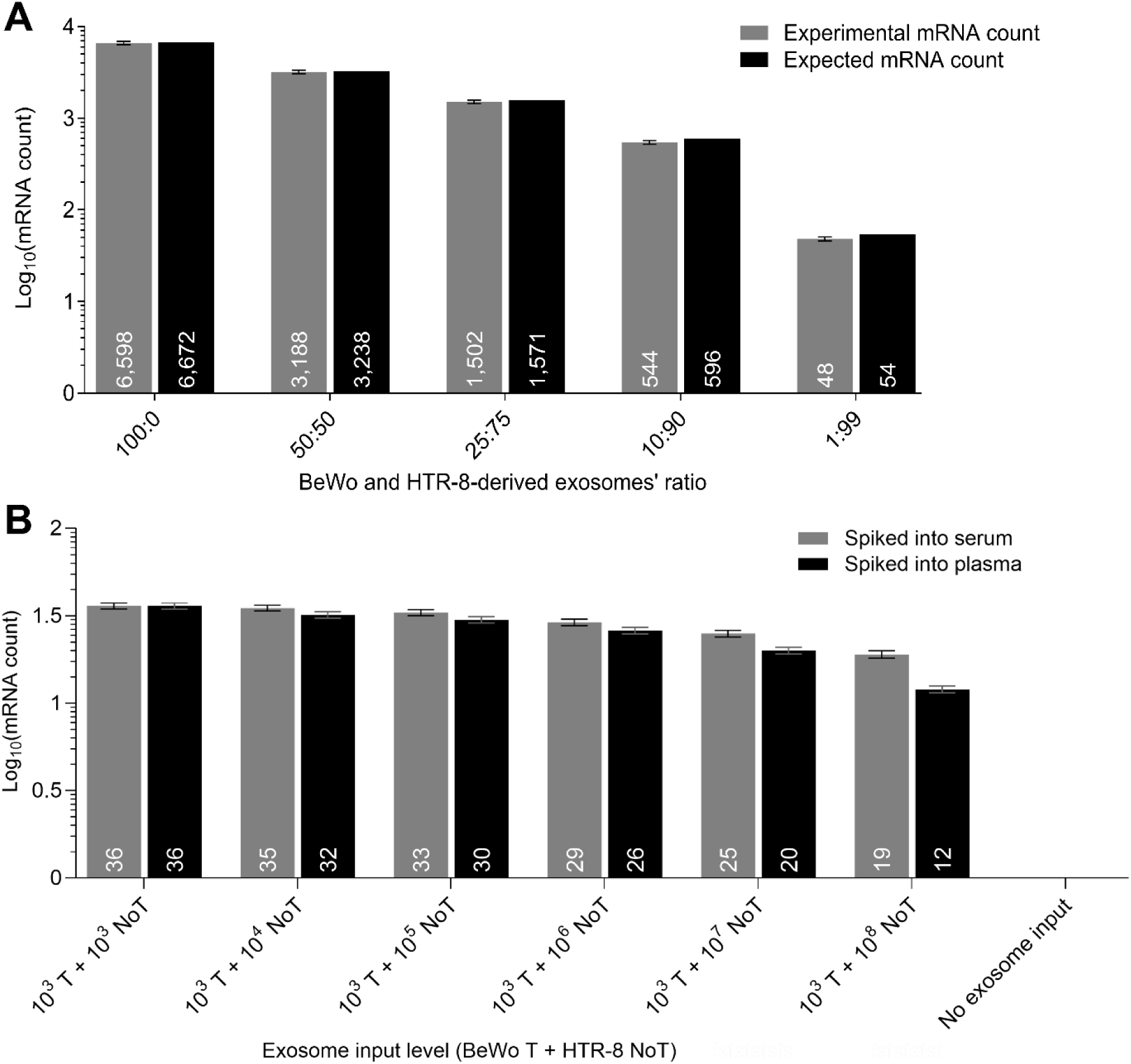
Exo-LAMP performance in heterogeneous exosome populations. (A) Exosome- depleted serum spiked with mixtures of PLAP^P^ BeWo-derived exosomes and PLAP^N^ HTR-8- derived exosomes at target-to-non-target ratios ranging from 100:0 to 1:99. Measured KISS1 mRNA copy numbers closely matched expected values across the dilution series, including at 1% target abundance. (B) Assay performance at the limit of quantification (LOQ) in heterogeneous serum and plasma matrices. PLAP^P^ BeWo-derived target (T) exosomes were fixed at 10^3^ exosomes per reaction and mixed with increasing amounts of PLAP^N^ HTR-8- derived non-target (NoT) exosomes ranging from 10^3^ to 10^8^ exosomes. Exo-LAMP generated detectable KISS1 mRNA signals in both exosome-depleted serum and plasma across all non-target backgrounds. Plasma spiked samples produced slightly lower mRNA counts than serum spiked, with the difference becoming more evident at higher non-target loads (10^7^ and 10^8^ exosomes). No detectable signal was observed in serum-only or plasma-only controls without spiked exosomes. Numbers within bars indicate estimated KISS1 mRNA copy counts for each condition.

To further evaluate assay robustness in a clinically relevant heterogeneous background, we fixed the BeWo-derived PLAP^P^ target exosome input at 10^3^ exosomes, corresponding to the LOQ of the Exo-LAMP assay, and mixed this with PLAP^N^ HTR-8-derived non-target exosomes ranging from 10^3^ to 10^8^ exosomes. The resulting mixtures were then spiked into serum and plasma matrices. As shown in Figure 6B, Exo-LAMP generated detectable KISS1 mRNA counts in both serum and plasma across the tested NoT exosome concentrations. Although the overall count pattern was comparable between the two matrices, plasma showed slightly lower mRNA counts than serum. This reduction became more pronounced at higher non-target exosome inputs, particularly at 10⁷ and 10⁸ exosomes. This decrease may reflect the greater biological complexity of plasma, where endogenous non-target exosomes and other matrix components could contribute to mild assay interference [42]. However, no background mRNA count was detected in serum-only or plasma-only controls without external exosome input, confirming that the measured signals came from the spiked target exosomes.

Together, these findings demonstrate that Exo-LAMP assay can reliably detect and semi- quantify low-abundance PLAP^P^ target exosomes at the LOQ level, even when they are present within highly heterogeneous matrices and greatly outnumbered by PLAP^N^ NoT exosomes.

## 4. CONCLUSION AND PERSPECTIVES

To the best of our knowledge, Exo-LAMP represents the first isothermal amplification-based strategy for exosomal mRNA analysis in an extraction free format that integrates selective EVs capture, nucleic acid release, and semi-quantitative detection within a single workflow. The assay combines 20 minutes of immunomagnetic enrichment of PLAP^P^ exosomes, 5 minutes of heat-mediated RNA release in the presence of an RNase inhibitor, and 25 to 30 minutes of RT-LAMP based detection of KISS1 mRNA, resulting in a total assay time of approximately 50 to 55 minutes. This streamlined sequence reduces the number of processing steps typically required for exosomal RNA analysis while maintaining selective enrichment of placenta derived EVs and compatibility with crude biological lysates. Collectively, these features support Exo-LAMP as a practical analytical framework that addresses key operational constraints associated with conventional exosomal mRNA workflows and enables more direct molecular access to placental signals.

Several aspects offer opportunities for further refinement to extend its analytical and translational scope. At present, amplification readout relies on a dedicated LAMP instrument, and the development of a fully portable or instrument free visual quantification module would further enhance its suitability for decentralized testing environments. Additional optimization of DNA-based primer design under crude lysate conditions may improve robustness against enzymatic interference and further stabilize amplification efficiency across diverse sample matrices. Integration of exosome capture, thermal lysis, and amplification into a closed microfluidic architecture with lyophilized reagents could further simplify handling and reduce the risk of contamination, while also improving reproducibility in resource limited settings. Finally, expanded evaluation using sequencing based benchmarking and clinically heterogeneous maternal plasma cohorts will be important to further define analytical performance, interindividual variability, and the broader applicability of circulating exosomal KISS1 mRNA as a biomarker of early placental dysfunction.

## ACKNOWLEDGEMENTS

M.A.S., F.H., M.T.T., and O.H.B.H. acknowledge the support of Higher Degree by Research (HDR) scholarships from Charles Sturt University.

