## Supplementary File for "Exo-LAMP: A Rapid Extraction-Free Isothermal Assay for Exosomal mRNA Detection"

*for*

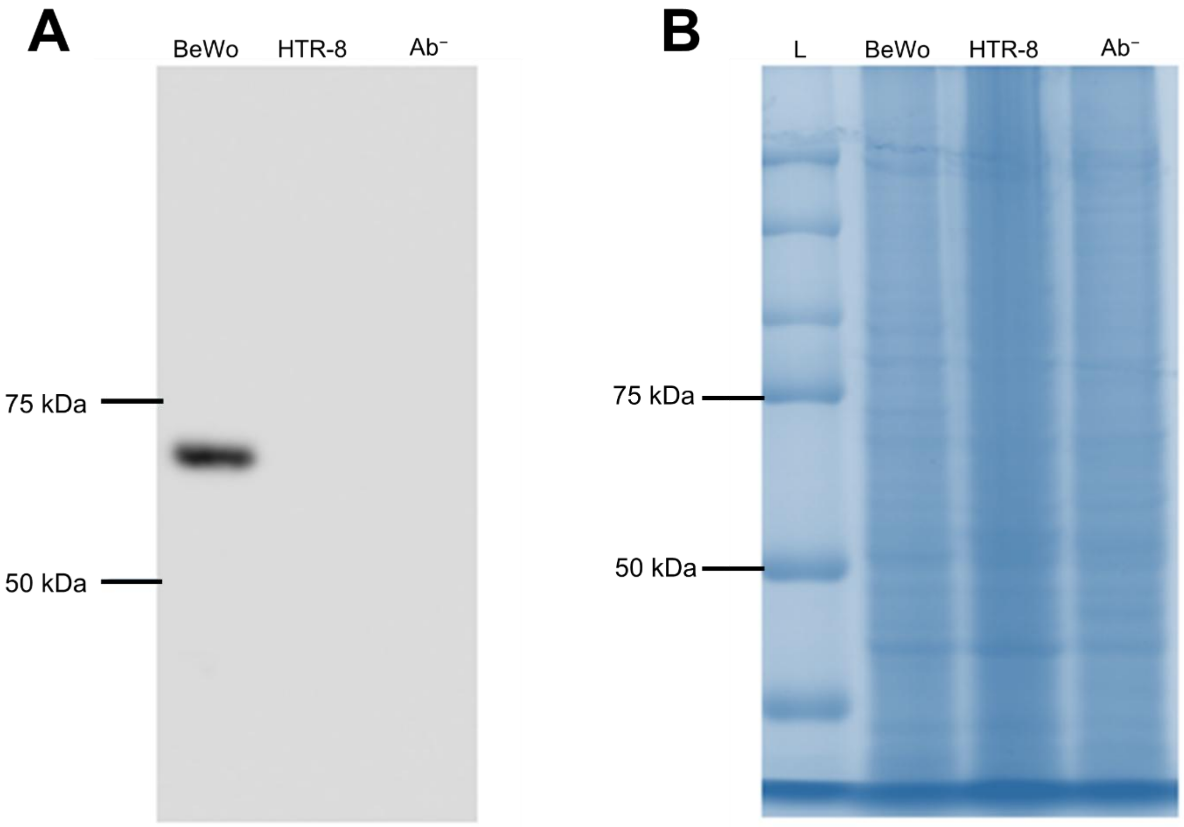


***Figure S1. Western blot analysis of placental alkaline phosphatase (PLAP) in exosomes.*** *(A) Immunoblot showing a distinct PLAP band in BeWo-derived exosomes, whereas no PLAP signal was detected in HTR-8-derived exosomes or in the anti-PLAP antibody-negative control ($Ab^{-}$), confirming selective PLAP expression in BeWo-derived exosomes. (B) Corresponding Coomassie blue–stained membrane showing total protein loading in each lane. L, protein ladder; $Ab^{-}$, anti-PLAP antibody-negative control.*

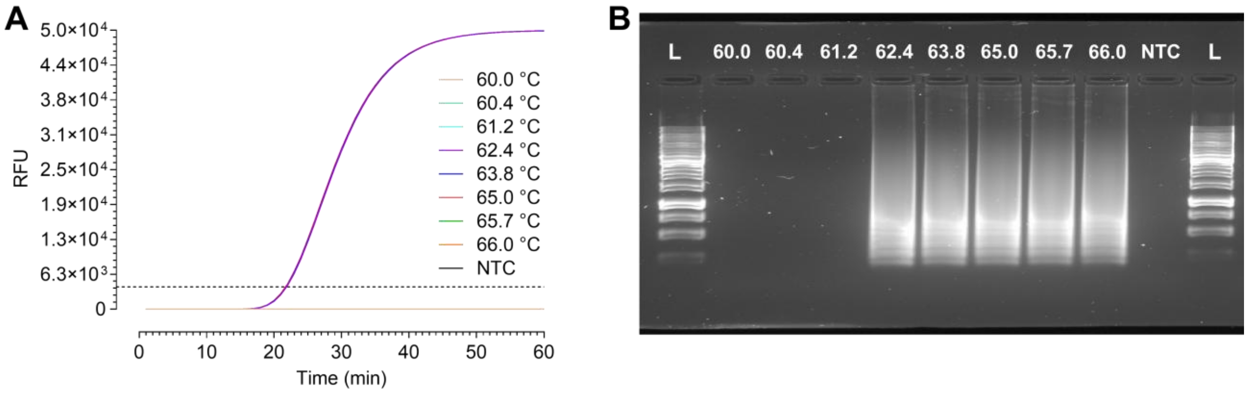


***Figure S2. Optimization of RT-LAMP reaction temperature.*** *(A) Real-time fluorescence profiles obtained from RT-LAMP reactions performed across a temperature gradient from 60.0 °C to 66.0 °C. Amplification was observed between 62.4 °C and 66.0 °C. (B) Agarose gel electrophoresis analysis of RT-LAMP products generated between 62.4 °C and 66.0 °C. L, ladder; NTC, no template control.*

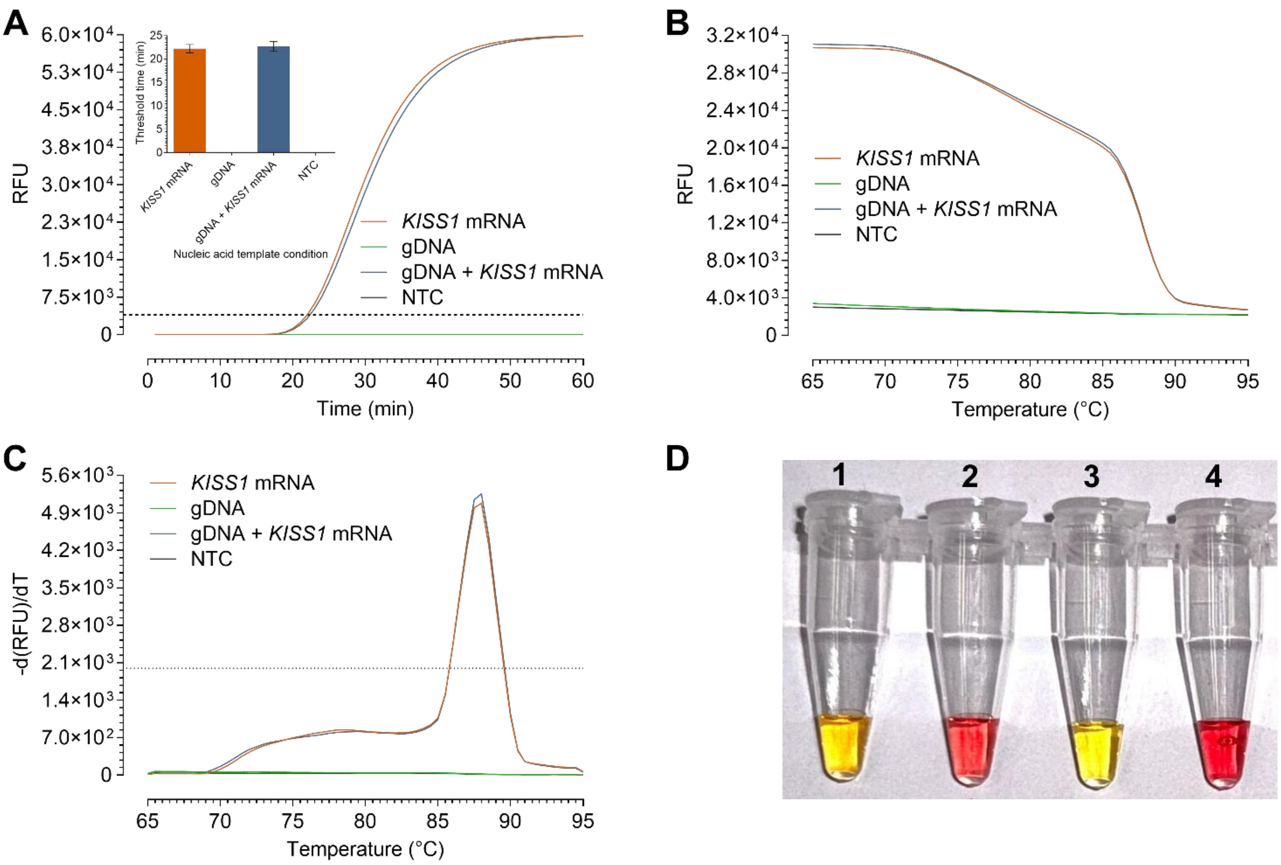


**Figure S3.** ***Evaluation of the LAMP primers specificity for KISS1 mRNA***. *(A) Real-time RT-LAMP amplification curves evaluating primer specificity for KISS1 mRNA. Strong amplification was observed in reactions containing KISS1 mRNA alone or KISS1 mRNA with genomic DNA (gDNA), whereas reactions containing only BeWo gDNA or the NTC showed no amplification. Inset: Corresponding threshold times derived from panel A. (B) Melt curve analysis of RT-LAMP products. (C) A single dominant melt peak was observed in reactions containing KISS1 mRNA alone or KISS1 mRNA with gDNA, confirming amplification of the same specific product. (D) End-point colorimetric readout of the specificity assay. Tubes containing KISS1 mRNA alone (Tube 1) or KISS1 mRNA with gDNA (Tube 3) changed from pink to yellow, whereas the gDNA-only (Tube 2) and NTC (Tube 4) reactions remained pink.*

**Optimization of exosome storage conditions**

To evaluate how freeze–thaw exposure under different storage conditions affects downstream exosomal RNA analysis, samples were preserved using four storage strategies, followed by exosomal *KISS1* mRNA quantification using the optimized immune magnetic isolation, heat lysis and RT-LAMP conditions. The tested conditions were: exosomes stored in serum at −20 °C, exosomes stored in serum at −80 °C, exosomes stored in serum containing dimethyl sulfoxide (DMSO) at −80 °C, and exosomes stored directly in their original CCM supplemented with DMSO at −80 °C.

When isolated exosomes were stored in serum at −20 °C (Figure S4A), the detected *KISS1* mRNA count remained relatively stable up to the first freeze–thaw cycle, after which a significant decline was observed compared with immediate analysis without freeze–thaw, as has also been reported previously for exosomal miRNA analysis [1]. A similar pattern was observed for storage in serum at −80 °C (Figure S4B). Although the *KISS1* mRNA count after the first freeze–thaw cycle was increased, the decline after subsequent cycles (Cycles 2 and 3) was more pronounced, and the counts eventually fell below those observed at −20 °C. This greater loss was due to freeze–thaw-induced disruption of isolated exosomes [2], which may increase free PLAP protein in the sample and thereby compromise subsequent MB@Ab-based capture of intact exosomes as well, resulting in lower recovery of amplifiable RNA. The addition of DMSO to serum improved exosomal preservation during storage at −80 °C, resulting in detectable *KISS1* mRNA counts that remained comparable to the non-freeze–thaw control up to the second freeze–thaw cycle (Figure S4C) [2]. The best preservation was observed when exosomes were stored within their source of origin, CCM supplemented with DMSO, without any prior isolation at −80 °C. Under this condition, *KISS1* mRNA counts remained comparable to the non-freeze–thaw control across all three freeze–thaw cycles (Figure S4D). This improved preservation in the original matrix is consistent with previous finding which showed that exosomal nucleic acid levels were maintained better when the exosomes were directly preserved in the source biofluid [3].

According to the *Minimal Information for Studies of Extracellular Vesicles 2023* (MISEV2023) recommendations, repeated freeze–thaw cycles should be minimized in EV studies because storage conditions can influence downstream measurements [4]. Consistent with this guidance, when storage was necessary in our workflow, exosomes were stored in DMSO-supplemented CCM at −80 °C and were used for no more than two freeze–thaw cycles.

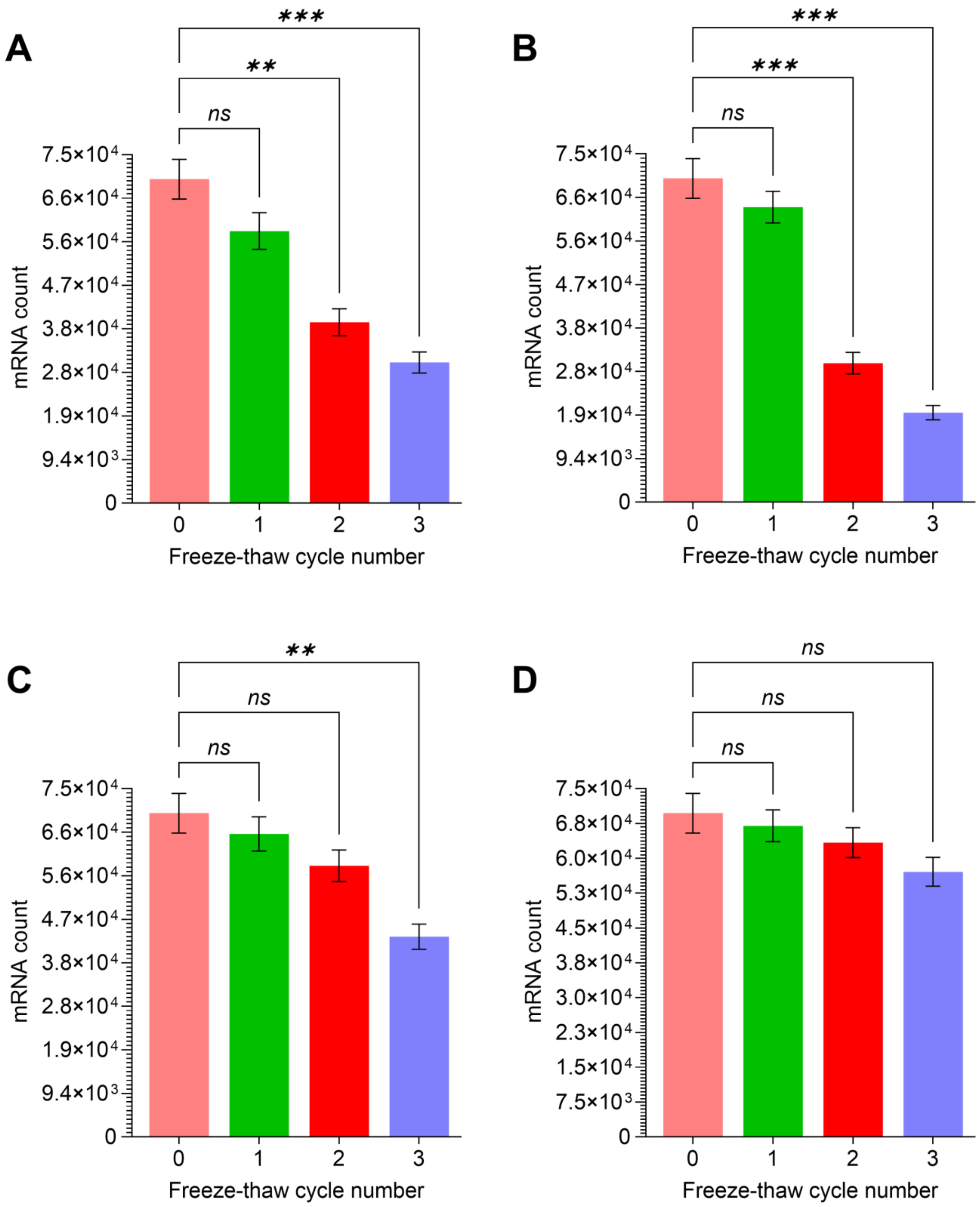


***Figure S4. Effect of freeze–thaw cycles under different storage conditions on detectable exosomal KISS1 mRNA.*** *Exosomes were stored under four conditions: in serum at -20 °C (A), in serum at -80 °C (B), in serum containing dimethyl sulfoxide (DMSO) at -80 °C (C), and in DMSO-supplemented conditioned culture medium (CCM) at -80 °C (D). 0 represents*

*immediate analysis without freeze-thaw, whereas 1, 2, and 3 represent samples subjected to one, two, and three freeze-thaw cycles, respectively.*

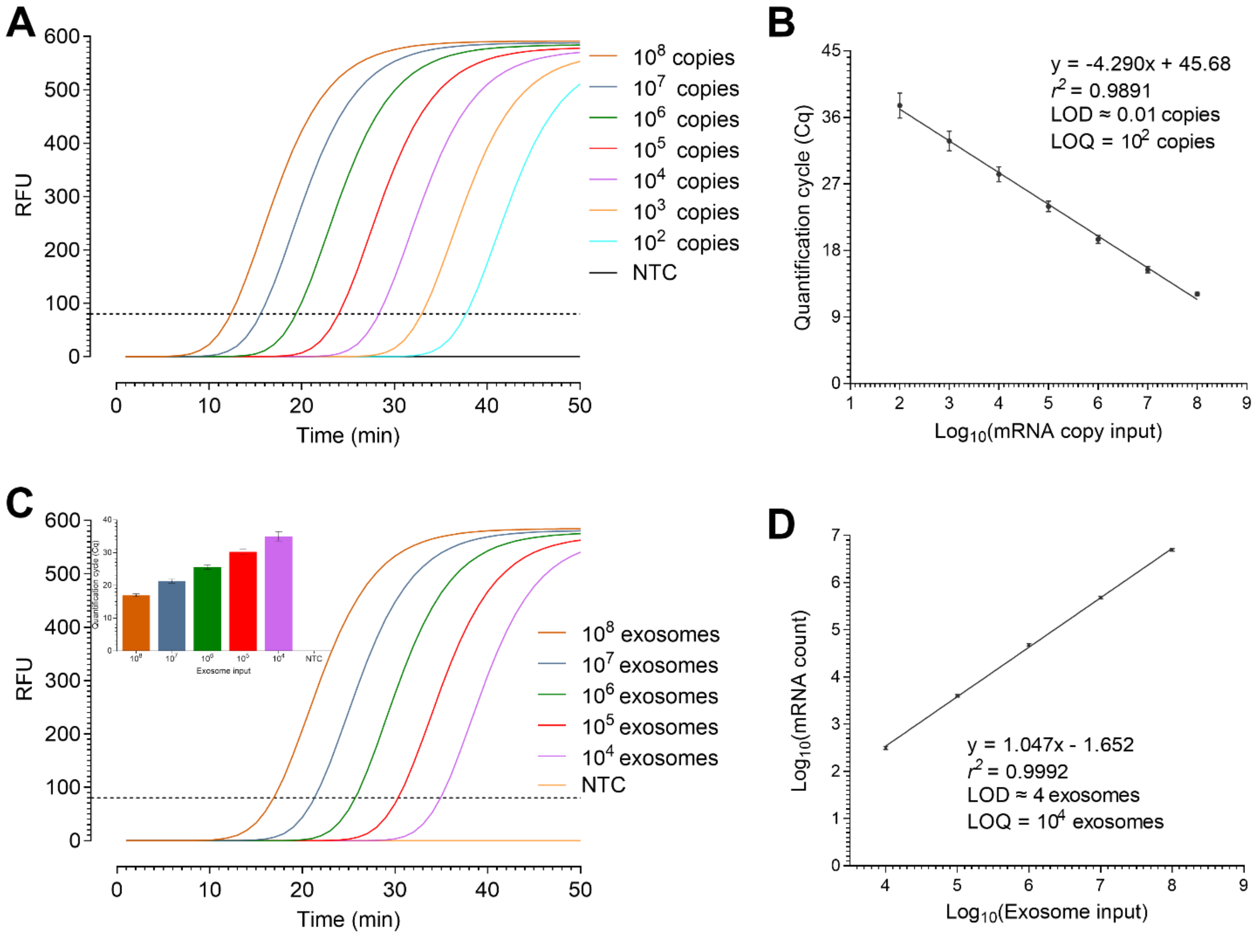


***Figure S5. Conventional RT-qPCR performance for KISS1 mRNA quantification.*** *(A) Real-time amplification curves from serial dilutions of synthetic KISS1 mRNA. (B) Standard curve generated using synthetic KISS1 mRNA. (C) Amplification curves from serial exosome inputs; inset shows the corresponding Cq values. (D) Standard curve showing the relationship between $Log_{10}$-transformed exosome input and estimated $Log_{10}$-transformed KISS1 mRNA copy number.*
